# The spatial network structure of intertidal meiofaunal communities derived from environmental DNA metabarcoding surveys in Northwest Iberia

**DOI:** 10.1101/2021.03.16.435605

**Authors:** Bruno Bellisario, Maria Fais, Sofia Duarte, Pedro E. Vieira, Carlos Canchaya, Filipe O. Costa

## Abstract

The identification of the patterns and mechanisms behind species’ distribution is one of the major challenges in ecology, having also important outcomes for the conservation and management of ecosystems. This is especially true for those components of biodiversity providing essential ecosystem functions and for which standard surveys may underestimate their real taxonomic diversity due to their high degree of cryptic diversity and inherent diagnosis difficulties, such as meiofaunal communities. Environmental DNA (eDNA) metabarcoding may provide a fast and reliable way to refine and scale-up the characterization of biological diversity in complex environmental samples, allowing to bypass such drawbacks and increase the resolution of biodiversity estimates. Moreover, the possibility of integrating eDNA metabarcoding-derived data with tools and methods rooted in network theory would deepen the knowledge of the structuring processes of ecological communities in ways that cannot be predicted from studying individual species/communities in isolation. Here, a sediment eDNA metabarcoding of mitochondrial cytochrome c oxidase I (COI) and the nuclear hypervariable V4 region of the 18S rDNA (18S) was used to reconstruct the spatial networks of intertidal meiofaunal OTUs from three estuaries of North-Western Iberian Peninsula. Null models were used to identify the role of environmental and spatial constraints on the structure of COI- and 18S-derived spatial networks and to characterize the macroecological features of surveyed phyla. Our results show the feasibility of eDNA metabarcoding, not only to capture a fair amount of diversity hard to detect with standard surveys procedures, but also to identify hierarchical spatial structures in intertidal meiofaunal assemblages. This suggests that exclusivity of occurrence rather than pervasiveness appears to be the norm in meiofaunal organisms and that niche-based processes predominantly drive the spatial aggregation and contemporary distribution of meiofaunal phyla within the system.

## **1.** Introduction

Advances in High-Throughput Sequencing (HTS) technologies have been allowing the characterization of biological diversity in complex environmental systems with unprecedented scale, speed and spatiotemporal density (Bik et al., 2012; Grey et al., 2018; Lacoursière-Roussel et al., 2018). In this context, the use of an environmental DNA (eDNA) metabarcoding approach may facilitate the identification of macroecological patterns and help solving fundamental ecological questions that could not be addressed by traditional biodiversity surveys (Drummond et al., 2015). eDNA-based biodiversity studies can also reveal more on the spatial partitioning of diversity and improve the knowledge on community assembly processes across the whole tree of life (Stat et al., 2017). The integration of *en masse* sequencing-derived data into macroecological frameworks can thus provide insights into the possible mechanisms underlying the structuring of communities, bypassing a series of issues related to diversity underestimation. Although not devoid of technical issues (Cristescu, 2014), eDNA metabarcoding increasingly represents a fast and reliable tool in biodiversity monitoring and conservation (Ji et al., 2013; Thomsen and Willerslev, 2015), providing also unprecedented opportunities to analyse the complexity of the relationships between species, and between species and their environment (Seymour et al., 2020a). One fruitful avenue of research is the integration of DNA-based approaches with network theory (Claire et al., 2019), where molecular data can be used to reconstruct species co-occurrence networks or even food webs (Compson et al., 2019; Djurhuus et al., 2020). However, data from eDNA metabarcoding surveys are seldom analysed by considering their spatial network configuration, where Operational Taxonomic Units (OTUs) can be linked to the sampling sites from where they have been surveyed to gain insight into the macroecological patterns and biogeographic distributions of organisms.

Spatial networks have interesting and universal properties that allow identifying recurrent structural patterns known to play a key role in the dynamic and stability of communities, as well as the spatial scale of influence of specific ecological processes determining the diversity and distribution of species (Fletcher Jr. et al., 2013; Gilarranz, 2020). For instance, modularity, that is, the tendency of a subgroup of sites and species to be linked to another subgroup of sites and species in a network, has proved to be a valuable tool to detect distinctive sets of sites (i.e., modules) with a greater number of common species (Borthagaray et al., 2018). The position of nodes within a modular network (i.e., their topological roles, *sensu* Guimerà and Amaral, 2005) can also provide useful information on the distributional range of species or the importance of sites/habitat patches for the interchange of species within the module and/or across the entire landscape (Carstensen et al., 2012, 2013; Borthagaray et al., 2014). This would allow identifying the mechanisms shaping the distribution of biodiversity, ultimately providing useful information for the delineation of management units (Borthagaray et al., 2018).

Species distributional patterns arise from a mix of different factors as, for instance, historical processes, dispersal limitation, intraspecific interactions and environmental constraints (Carstensen et al., 2013; Pigot and Tobias, 2013; Borregaard et al., 2016). The identification of the spatial boundaries at which abiotic and biotic processes determine the mechanisms of local and regional assemblage (e.g., competition, environmental filtering, dispersal) potentially allow for high-resolution assessments of biodiversity in different ecosystems and implementation of focused conservation strategies (Borthagaray et al., 2018). This is paramount for poorly known layers of biodiversity providing essential functions in threatened ecosystems (Solan et al., 2004), and for which standard surveys may underestimate the real taxonomic diversity due to the high degree of cryptic diversity and inherent diagnosis difficulties (Lobo et al. 2017).

Recent studies have shown the feasibility of eDNA metabarcoding to recover a greater amount of diversity in estuarine benthic communities when compared with morphological- only methods, slightly exposing differences in species composition among natural communities and providing high quality and auditable species identifications (Lobo et al., 2017; Steyaert et al., 2020). Within the benthic domain, meiobenthos represents a fundamental component characterized by a limited reproductive output and dispersal, as well as by rapid generational turnovers, playing a key role in regulating important ecological, trophic and sedimentary processes (Fenchel and Finlay, 2004; Giere, 2009; Schratzberger and Ingels, 2018). Traditionally comprising organisms between 0.63 µm and 1 mm in size, meiofauna is actually represented by a dense community of microeukaryotes able to adapt physiologically and anatomically to living in the interstitial sedimentary environment and to respond quickly to environmental changes (Kennedy and Jacoby, 1999). Although their short life cycles, reproductive K-strategies and lack of pelagic larvae would suggest comparatively narrow-species distributions, many meiobenthic species are considered to be amphioceanic or even cosmopolitan (e.g., the so-called meiofauna paradox, Giere, 2009 and references therein). Indeed, the fast rising of molecular technologies with improved detection accuracy has led to the reappraisal of the widespread or cosmopolitan status of some species complexes (Fontaneto, 2019). In particular, several genetically divergent species with a limited geographical distribution were discovered through the use of DNA-based approaches (Guardiola et al., 2016; Fonseca et al., 2017; Mills et al., 2017; Faria et al., 2018; Worsaae et al., 2019). Moreover, evidence on the patchy distribution of meiofauna has been found through the application of DNA metabarcoding and ecological network analysis, highlighting high levels of both *α* and *β*-diversity (Fais et al., 2020b).

The diversity and distribution of meiofaunal assemblages are influenced by the response of species to both large-scale gradients (e.g., temperature, salinity) and local variation in physical-chemical variables (e.g., tidal exposure, organic matter). However, despite recent findings suggest the prevalence of niche-based mechanisms in structuring meiofaunal communities, the spatial extent at which community assembly processes act is not yet clear. Large scale studies and adequate analytical frameworks able to investigate meiofaunal assemblages as a whole are required (Gansfort et al., 2020).

In this paper, we coupled eDNA metabarcoding with network theory to disentangle the role of geographic and environmental constraints in determining the distributional patterns of intertidal meiofaunal communities sampled in three estuaries along the North-Western Iberian Peninsula. Two molecular markers, the mitochondrial cytochrome c oxidase I (COI) and the hypervariable V4 region of the 18S rDNA (18S), have been employed in standardized survey campaigns in three estuaries with different geomorphological and environmental characteristics. Null models were used to test specific hypotheses relating environmental and spatial constraints on the observed modular partitioning of eDNA-derived spatial networks, and to characterize the macroecological features of surveyed phyla by means of their local and regional distributional range.

## 2. Materials and Methods

### 2.1 Study area

The study area covered approx. 300 km of distance in the North-Western Iberian Peninsula, between Galicia (Spain) and the North-Central regions of Portugal (Fig.1). Four sampling stations within three estuarine systems, characterized by different geomorphological and environmental characteristics, were selected for metabarcoding intertidal meiofauna (Table 1): i) south margin of Ría de Vigo, ii) Lima estuary and, iii) Ría de Aveiro.

**Figure 1.**
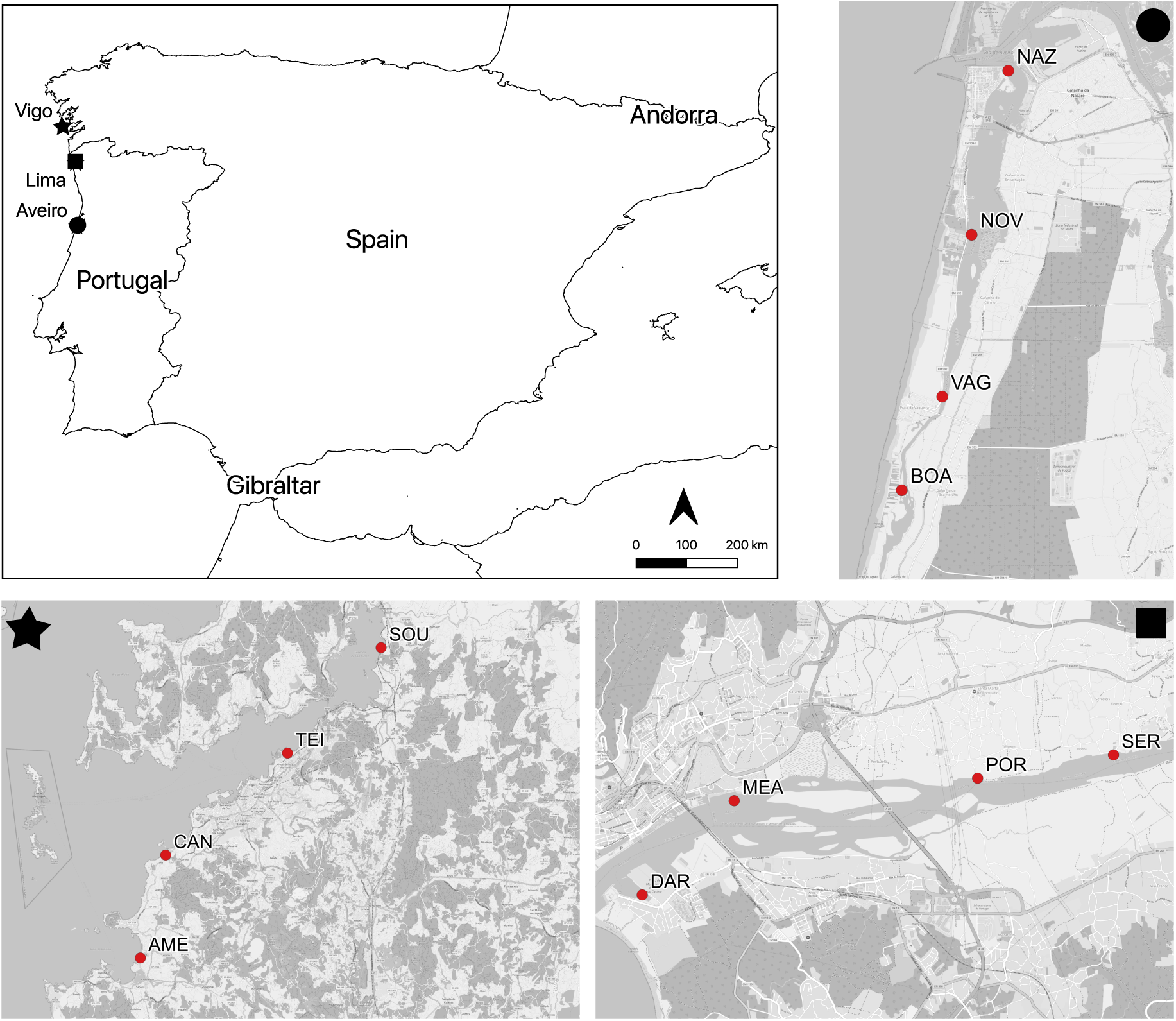
– Geographic scope, estuary affiliation and locations of sampling stations. See Table 1 for acronyms of sampling stations.

**Table 1.** – Environmental and geographic characterization of sampling stations. Values are reported as mean ± SD. S: salinity; SS: sandy sediments; MS: muddy sediments; TOM: total organic matter. Sampling stations acronyms: DAR, Darque; MEA, Meadela; POR, Santa Marta de Portuzelo; SER, Serreleis; NAZ, Gafanha da Nazaré; NOV, Costa Nova; VAG, Vagueira; BOA, Boa Hora; AME, Praia America; CAN, Canido; TEI, Enseada de Teis; SOU, Soutoxuste.

|  | Estuary | Latitude | Longitude | S* (psu) | SS (g) | MS (g) | TOM (%) |
| --- | --- | --- | --- | --- | --- | --- | --- |
| DAR | Lima | 41.683 | -8.828 | 19.56 | 87.125 (± 9.426) | 10.345 (± 2.737) | 0.0135 (± 0.001) |
| MEA | Lima | 41.695 | -8.812 | 5.66 | 90.755 (± 17.883) | 2.505 (± 0.064) | 0.057 (± 0.003) |
| POR | Lima | 41.698 | -8.772 | 1.78 | 131.88 (± 51.746) | 6.825 (± 1.690) | 0.0695 (± 0.006) |
| SER | Lima | 41.701 | -8.749 | 2.12 | 115.105 (± 6.725) | 4.205 (± 0.460) | 0.035 (± 0.013) |
| NAZ | Aveiro | 40.643 | -8.736 | 30.3 | 105.065 (± 5.650) | 19.83 (± 0.877) | 0.045 (± 0.007) |
| NOV | Aveiro | 40.603 | -8.747 | 24.28 | 113.445 (± 12.297) | 17.49 (± 1.047) | 0.015 (± 0.007) |
| VAG | Aveiro | 40.565 | -8.757 | 21.09 | 82.33 (± 0.226) | 17.325 (± 3.076) | 0.025 (± 0.007) |
| BOA | Aveiro | 40.542 | -8.769 | 18.52 | 64.505 (± 1.619) | 19.96 (± 0.806) | 0.04 (± 0.001) |
| AME | Vigo | 42.129 | -8.819 | 31.5 | 76.78 (± 4.653) | 4.28 (± 1.216) | 0.001 (± 0.001) |
| CAN | Vigo | 42.194 | -8.798 | 31.65 | 65.94 (± 0.820) | 10.725 (± 2.906) | 0.005 (± 0.001) |
| TEI | Vigo | 42.258 | -8.695 | 31.2 | 54.345 (± 3.882) | 37.21 (± 0.438) | 0.013 (± 0.001) |
| SOU | Vigo | 42.324 | -8.615 | 29.3 | 39.455 (± 2.595) | 48.965 (± 4.575) | 0.021 (± 0.001) |

\*Values are reported as absolute mean without SD due to the null variability of salinity values between replicates.

Ría de Vigo (Fig. 1) is a V-shaped estuarine basin classified as a fjord (Pritchard, 1967), belonging to the coastal system of the Galician Rías Baixas. This basin is characterized by an external area influenced exclusively by ocean/tidal currents, and an innermost one influenced by both river flow and ocean/tidal currents (García-Moreiras et al., 2018). Upwelling events are typical during the warmer months and anthropogenic impacts are high due to coastal industrialization, massive navigation, dredging, wastewater inputs, among other disturbances (García-Moreiras et al., 2018). Lima estuary (Fig. 1) is a drowned river valley (Pritchard, 1967), characterized by mesotidal condition with an evident saline wedge during tidal flood, being a moderately disturbed area due to agriculture, industrial and dredging activities (Costa-Dias et al., 2010). Ría de Aveiro (Fig. 1) is a coastal lagoon or bar-built estuary (Pritchard, 1967), connected directly to the ocean through an inlet and dominated mainly by tidal forces. The area is also characterized by marked anthropogenic impacts due to the presence of industrial, agricultural, aquaculture and navigation activities (Bueno-Pardo et al., 2018).

### 2.2 Sampling strategy and sediment analysis

Three samples of intertidal sediment were collected in each station of each estuarine system (3 samples x 4 stations x 3 estuaries), during June (Lima estuary and Ría de Aveiro) and July 2018 (Ría de Vigo). The first 5 cm (± 0.5 cm) of sediment were collected directly to 50 mL sterile Falcon^®^ tubes and stored immediately in dry ice and later at -20 °C, until eDNA extraction. At each station, surface water salinity was measured through a Multiparameter Sea Gauge YSI EXO 2, and additional sediment samples were collected to perform granulometric and TOM analyses. Sediment samples for particle size measurement were first dried at 50 °C and then subjected to mechanical sifting in order to be classified according to the Udden-Wentworth scale (Udden, 1914; Wentworth, 1922). Granules between 0.125 and 2 mm were grouped as sandy sediments, while those below 0.125, as muddy sediments. To calculate the TOM, the fine fraction of the sediments (< 0.0063 mm) was burned in a muffle at 400 °C for 4 hours and subsequently weighted to determine the difference compared to the initial weight (more details can be found in Fais et al., 2020a).

### 2.3 Molecular analyses

eDNA was extracted directly from 10 g (± 0.5 g) of sediments to avoid the loss of individuals with soft and/or small body mass, using the QIAGEN^®^ PowerMax Soil DNA Isolation kit (cat#12988-10), following optimized procedures from a previous work on meiofauna (Fais et al., 2020a). Sediment-free negative extraction controls were used to screen out any contamination of the solutions of the DNA extraction kits and labware used. Sub-regions with 313 bp of the gene coding for the mitochondrial cytochrome c oxidase I (COI) and ∼400 bp of the hypervariable V4 region of the 18S rDNA (18S) were PCR amplified with the mICOIintF/LOBOR1 (Forward : 5’- GGWACWGGWTGAACWGTWTAYCCYCC - 3’ / Reverse: 5’- TAAACYTCWGGRTGWCCRAARAAYCA- 3’; Leray et al., 2013 and Lobo et al., 2013, respectively) and the TAReuk454FWD1/TAReukREV3 (Forward: 5’ - CCAGCASCYGCGGTAATTCC - 3’/ Reverse: 5’ - ACTTTCGTTCTTGATYRA - 3’; Lejzerowicz et al., 2015) primer pairs, respectively. A total of 72 amplicon libraries were submitted to high-throughput sequencing in an Illumina MiSeq platform at Genoinseq (Biocant, Cantanhede, Portugal). PCR reactions were performed using 0.3 μM KAPA HiFi HotStart PCR Kit, using 12.5 ng of DNA template in a total 25 μL volume for each amplicon. The PCR conditions were: 3 min denaturation at 95 °C; 35 cycles at 98 °C for 20 s, 60 °C for 30 s and 72 °C for 30 s; a final extension at 72 °C for 5 min (COI); and 3 min denaturation at 95 °C; 10 cycles of 98 °C for 20 s, 57 °C for 30 s and 72 °C for 30 s; 25 cycles of 98 °C for 20 s, 47 °C for 30 s and 72 °C for 30s; a final extension at 72 °C for 5 min (18S). Negative PCR controls were included in all amplification reactions.

### 2.4 Bioinformatic procedures

Good quality R1 and R2 reads (> 150 bp and > Q25 in a window of 5 bp; Schmieder and Edwards, 2011) without adapters were used to produce contigs sequences for further processing, by following and adapting operational procedures in mothur v.1.39.5 (Schloss et al., 2009; Kozich et al., 2013). OTU clustering was performed at 97% of similarity in order to best represent the phylogenetically diverse meiofaunal community and to seek a reliable compromise for both molecular markers (Botnen et al., 2018). OTUs with < 9 sequences (rare OTUs) and present only once in the COI-A or V4 datasets (singleton OTUs) were removed to elude sequencing artefacts and cross-talk errors on diversity estimation (Leray and Knowlton, 2017; Edgar and Flyvbjerg, 2018). Sequences within the 3% threshold for each OTU centroid were considered as representative (Rognes et al., 2016) and used to perform BLAST (Basic Local Alignment Search Tool v.2.6.0; Benson et al., 2018; last access: April 2019) against the non-redundant nucleotide (nt) GenBank^®^ library with the following criteria: -evalue 1e^-30^, -max_target_seqs 50, -perc_identity 70 (executed in GNU Parallel, Tange, 2011). The top 50 BLAST matches were analyzed by the MEtaGenome ANalyzer (MEGAN v.6.13.1; Huson et al., 2016), assigning and correcting the nomenclature to the most certain taxonomic level in accordance with WoRMS (World Register of Marine Species, last access: April 2019) and NCBI (National Centre for Biotechnology Information, last access: April 2019) databases. Only OTUs attributed to the temporary and/or permanent meiofauna were considered to design the OTU tables employed in this study. Raw sequences without barcodes/adapters are available on the Sequence Read Archive (SRA) of NCBI, in the BioProject PRJNA611064, under the following accession numbers: SAMN14944165 (COI - Lima Estuary); SAMN14944245 (18S – Lima Estuary); SAMN14967338 (COI – Ría de Vigo); SAMN14967349 (18S – Ría de Vigo); SAMN14967353 (COI – Ria de Aveiro); and SAMN14967358 (18S – Ria de Aveiro).

### 2.5 Data analysis

OTU richness was considered for each station of three estuarine systems. Phylum and order levels were considered for the qualitative evaluation of the meiofauna composition along the stations of each estuary. Venn diagrams were produced in mothur and re-designed appropriately in R v. 4.0.2 (R Development Core Team, 2019) with the *VennDiagram* package (Chenn and Boutros, 2011). Both the percentage of OTU replacement and unique OTUs were subsequently extrapolated. Hence, shared OTUs were assessed for consideration of their exclusivity/pervasiveness: ‘very exclusive’ OTUs (i.e., OTUs present only in one estuary); ‘partially exclusive’ OTUs (i.e., OTUs shared between two estuaries); and ‘pervasive’ OTUs (i.e., OTUs occurring in the three estuaries).

### 2.6 Spatial network construction

The spatial network structure of meiofaunal communities was modelled as a bipartite network, where links have been established between two sets of nodes (OTUs and sampling stations), but not between nodes of the same set. Starting from the raw sequence data and for each primer used (COI and 18S), we compiled the incidence matrix whose values expressed the total reads of a given OTU (row) in a given sampling station (column). OTU tables were optimized to avoid: i) possible biases related to unequal sequencing effort among samples and artifacts introduced in PCR and sequencing processes and, ii) interspecific variation in the intra-genomic copy number of barcoding regions (Toju, 2015). Sequencing reads were then rarefied to a subsampling size corresponding to the smallest number of total reads in the original sample-level matrix to remove extremely rare reads. Matrices were finally converted into a bipartite graph, in which a single link was created for every non-zero element.

### 2.7 Modularity

Here, we focused on modularity (*Q*), a particular network organization where highly linked subgroups of nodes constitute modules, and a few nodes connect modules together to form one large coherent network. From an ecological point of view, modularity provides a formal description of the pattern of aggregation between species, populations, communities or habitat patches, with modules defining the spatial limits at which specific ecological and evolutionary processes occur (Fletcher Jr et al., 2013; Gilaranz 2020).

Modularity was measured by means of the LP-BRIM algorithm available in the R package *lpbrim* (Poisot and Stouffer, 2015), due to its precision and speed in identifying meaningful community structures in large-scale bipartite networks. LP-BRIM is based on two well-known algorithms, LP (label propagation, Raghavan et al., 2007), a very fast community detection algorithm and BRIM (Bipartite Recursively Induced Modules, Barber, 2007), an algorithm that applies a recursively induced division between the two types of nodes in bipartite networks to generate better community structures.

### 2.8 Macroecological classification of OTUs

We classified the distributional patterns of OTUs by means of their topological linkage in modular networks, that is, the extent to which individual OTUs were connected both within each module and across modules. For each OTU in each spatial network, we calculated the connectivity (*z*) and participation (*p*) coefficients, two measures able to provide useful information on the distributional range size of nodes in a network (Guimerà and Amaral, 2005). *z* measures how well-connected a node is to other nodes in its own module, while *p* measures how well-distributed links of a node are among different modules. Nodes can thus be assigned to different ‘topological roles’ on the basis of their position in the bidimensional space given by conventionally defined thresholds in the values of *z* and *p* (Guimerà and Amaral, 2005). For instance, nodes having a distributional range restricted to few sampling localities within the same module can be defined as ‘peripherals’ (i.e., *p* < 0.62 and *z* < 2.5), while those having a widespread distribution encompassing different modules as ‘hubs’ (i.e., *p* > 0.62 and *z* > 2.5) (more information can be found in Guimerà and Amaral, 2005).

However, since the quantification of the role alone could be uninformative without assessing the taxonomic rank to which OTUs belong, we performed a Factor Analysis of Mixed Data (FAMD) to identify the distributional range size of different phyla in the networks. FAMD is a principal component method particularly useful for dealing with both continuous and categorical variables by combining a principal component analysis (PCA) for continuous variables and a multiple correspondence analysis (MCA) for categorical variables (Pagès, 2004). FAMD was thus applied to both COI and 18S OTUs lists with phyla, *z* and *p* as variables, by using the package *FactoMiner* of R (Lê et al., 2008).

### 2.9 Null model analyses

We used a null model approach to test for the influence of spatial and geographic constraints on the observed modular structure of intertidal meiofaunal assemblages. Specifically, we employed a novel class of null models, namely correlation-informed, which combines classical approaches (e.g., the swap algorithm, Gotelli and Entsminger, 2001) with tools from community ecology (e.g., joint statistical modelling, Mora et al., 2019). Basically, a correlation-informed null model allows assessing the predictive power of a given correlation matrix defining the relationship among the *m* columns (or *n* rows) on the structural pattern observed within an incidence matrix. Predictions are then defined by fitting the observed links to a logistic regression using generalized linear mixed models (Mora et al., 2019). The estimated rewiring probability (*p^ij^*) represents a bias in the null model, providing a way of weighting the randomization process based on the correlation matrix.

Different informed null models have been employed to quantify the influence of environment, geography and estuary affiliation (predictors) on the pattern of OTU distribution measured by modularity. For each predictor, we derived the corresponding Gaussian correlation structure as, *V*_cov_ = (1 - *N*) exp [(*D*/*D*_max_)^2^], where *D* is a generic distance matrix and *N* is a matrix having diagonal elements zero and off-diagonal elements *n^ij^* = *η*, a parameter used to avoid perfectly correlated off-diagonal elements when generating correlation structures. Here we followed Mora et al. (2019) and set *η* = 0.01. As environmental filtering and dispersal processes are known to be scale dependent (Kinlan et al., 2005; Menegotto et al., 2019), we used the geographic distance and estuary affiliation to generate a class of correlation structures by taking into account their role in determining the correlation between observations (see Table 2 for a complete list of correlation structures used in this study).

**Table 2.**
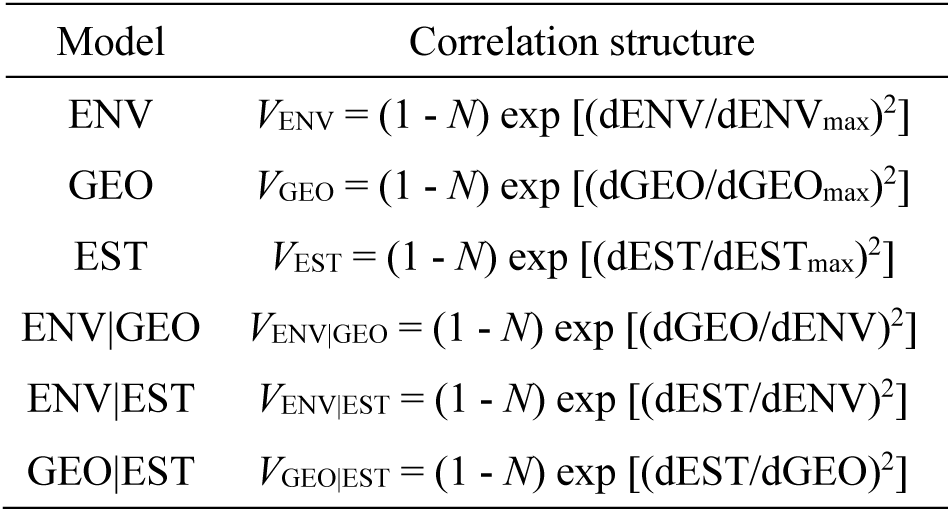
– List of Gaussian correlation structures employed in the correlation-informed null model analysis. dENV, dGEO and dEST are the environmental, geographic and estuary affiliation distance matrices, respectively. The symbol | identifies the class of correlation structures taking into account the role of geography (dGEO) and estuary affiliation (dEST) in determining the correlation between observations. *N* is a matrix having diagonal elements zero and off-diagonal elements *n_ij_* = *η*, corresponding to the nugget effect avoiding perfectly correlated off-diagonal elements in correlation structures (here *η* = 0.01, Mora et al. 2019).

To compute the environmental distance matrix between sampling stations, we first performed a PCA on the measured environmental parameters (Table 1) to reduce dimensionality and avoid collinearity among variables. The first two axes explained a cumulative percentage of variance of ca. 88% (PCA1 = 67.581% and PCA2 = 20.475%) and were used to calculate the Euclidean distance matrix. Pairwise geographic distances between sampling stations were calculated with a least-cost path approach, by using land areas (masked using the European Environment Agency coastline polygon 1:100000) as a barrier in distance calculations. This method allows for a more realistic evaluation of the effective spatial separation than the more commonly used Great Circle or Euclidean distances due to the complex geometry of coastlines in the study area (Rattray et al., 2016). We also considered the estuary affiliation to account for the estuary-level effects in determining the correlation structure among sampling stations. We therefore assigned to each station a numeric code corresponding to the estuary it belonged (Table 1 and Fig. 1) and then calculated the euclidean distance matrix. Finally, we tested for randomness in the observed modularity by means of an uninformed null model using a fixed-fixed algorithm, whereby elements in the incidence matrix were randomized fixing both the number of OTUs per site and the relative frequency of appearance of each OTU.

To test whether the modular structure observed in the empirical incidence matrices was significantly non-random compared to the data generated by the uninformed and informed null models, we used the *z*-score, *z* = (*Q*_obs_ - *μ_Q_*) / *σ_Q_*, where *Q*_obs_ is the observed modularity and *μ_Q_* and *σ_Q_* the mean and standard deviation of 100 randomized matrices, respectively. Values of *z* between -1.96 and 1.96 indicate a not significant effect of a given predictor, while values greater than 1.96 and lower than -1.96 indicated a significant over and under representation of the observed pattern. Null model analyses were performed in R by using the package *RESOLDRE* (Mora et al., 2019). Data and codes written for this manuscript will be available upon request.

## 3. Results

### 3.1 Environmental characterization of sampling areas

Sampling areas showed a clear subdivision along the first axis of the PCA (Fig. 2), with all sampling stations within the Lima estuary characterized by the highest values of total organic matter and sandy sediments, while those in Ría de Vigo by the highest values of muddy sediments and salinity (Fig. 2). Stations in Ría de Aveiro were characterized by intermediate values of analysed parameters and a quite lower environmental variability as in Lima estuary (Fig. 2). Conversely, Ría de Vigo displayed a marked subdivision of sampling stations along the second axis of the PCA, with the innermost ones (SOU and TEI, Fig. 2) characterized by low salinity and high percentage of muddy sediments and the most external (CAN and AME, Fig. 2) by an opposite trend.

**Figure 2.**
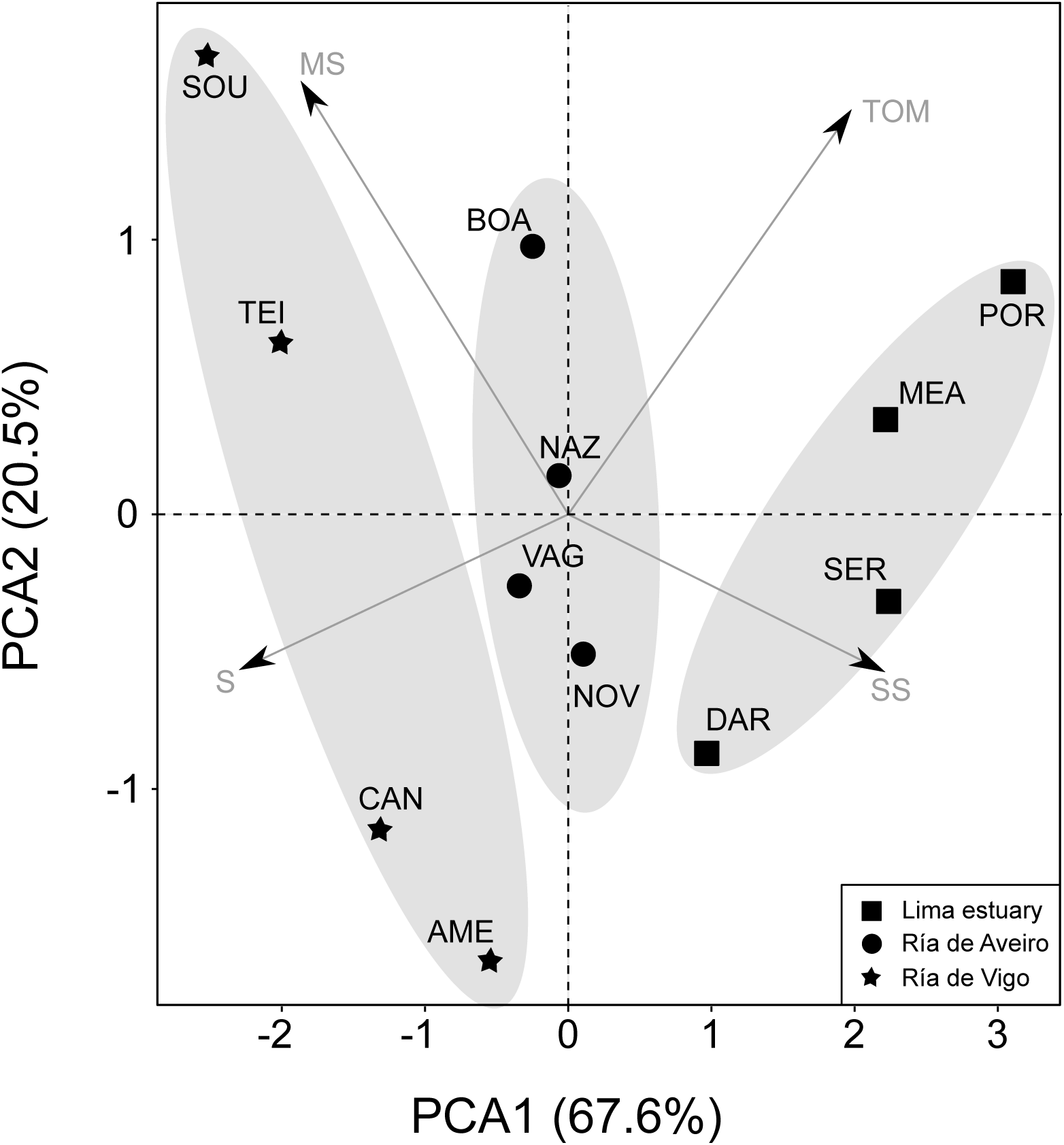
– Multivariate ordination pattern of sampling stations given by the Principal Component Analysis. MS: muddy sediments; TOM: total organic matter; SS: sandy sediments; S: salinity. See Table 1 for acronyms of sampling stations and estuary affiliation.

### 3.2 OTU diversity patterns

The number of initial sequences following the combination of the paired-end reads was 1,765,470 (COI) and 1,101,851 (18S). Subsequent steps of de-multiplexing and filtering reduced the number to 611,639 and 463,167 for COI and 18S, respectively. The latter corresponded to 789 (COI) and 712 (18S) OTUs, of which 8 (COI) and 21 (18S) did not have any taxonomic attribution. OTUs ascribed to meiofauna were 129 for COI and 143 for 18S, with a predominance of Animalia, Chromista and Protozoa taxa, for both primer pairs. For each region and primer pair, the richest stations in OTUs were SOU for Vigo, NOV for Aveiro and POR for Lima (Fig. S1 in Supplementary Materials). The 18S recovered greater OTU richness than COI, except for Vigo stations; but the COI primer pair recovered, on average, a higher number of phyla than 18S (Fig. S2 in Supplementary Materials). Qualitatively, both primer pairs were able to identify, preferentially or exclusively, certain groups of meiofauna in different taxonomic ranks and for each examined estuary (Fig. S2 in Supplementary Materials).

Patterns of OTU’s distribution showed a high turnover within and between the stations of each estuarine system, as well as between the three estuaries. In particular, we found high values of within-station turnover (80-92% for COI and 77-92% for 18S) with lower values for Ría de Vigo (both COI and 18S) and higher for Ría de Aveiro (COI) or Lima (18S) (Fig. S3 in Supplementary Materials). The overall turnover between estuaries was approximately 89% for COI and 83% for 18S (Fig. S3 in Supplementary Materials). The Ría de Vigo - Lima estuary pair had the highest values for both primer pairs (∼ 96%), while the Ría de Vigo - Ría de Aveiro and Lima estuary - Ría de Aveiro pairs displayed, on average, the lowest values for COI and 18S (∼ 83% and ∼ 74%, respectively). Furthermore, Ría de Vigo had the highest number of exclusive OTUs for both primer pairs compared to other two estuaries.

The within-estuary turnover was higher than within-stations, with approximately 96% for COI and 94% for 18S, highlighting the absence of shared OTUs among some stations within the same estuary (Fig. S3 in Supplementary Materials). Overall, OTU exclusivity was greater than OTU pervasiveness for both primer pairs, with ∼ 33% of OTUs surveyed through COI primer classified as ‘very exclusive’, ∼ 29% as ‘partially exclusive’ and only ∼ 10% as ‘pervasive’, while for 18S ∼ 29% were classified as ‘very exclusive’ and ∼ 31% as ‘partially exclusive’. OTU pervasiveness, or widespread occurrence, was found to be uncommon among assessed communities regardless of the primer pair, with only 12 (∼ 10%) and 24 (∼ 17%) found in all the three estuaries for a total of 129 and 143 detected in the study using COI or 18S, respectively (Fig. S3 in Supplementary Materials).

### 3.3 Spatial network structure

Modularity showed the partition of the COI-metabarcoding-derived spatial network in seven modules with a modularity value of *Q* = 0.465. Interestingly, all sampling stations in Lima estuary belonged to the same module (Fig. 3), while stations located in Ría de Aveiro were subdivided in two distinct modules, and those of Ría de Vigo formed four distinct and well-defined modules (Fig. 3). Results showed key differences between null models, especially when accounting for the spatial and estuary dependence on environmental and geographic distances. This class of informed null models showed the same pattern of overrepresentation as that observed when using the uninformed null model (Fig. 3). However, differences in environmental conditions and dispersal processes acting at finer scale appears to be particularly informative for determining the modular structure, since data generated by null models are much better at reproducing the empirical modularity (Fig. 3).

**Figure 3.**
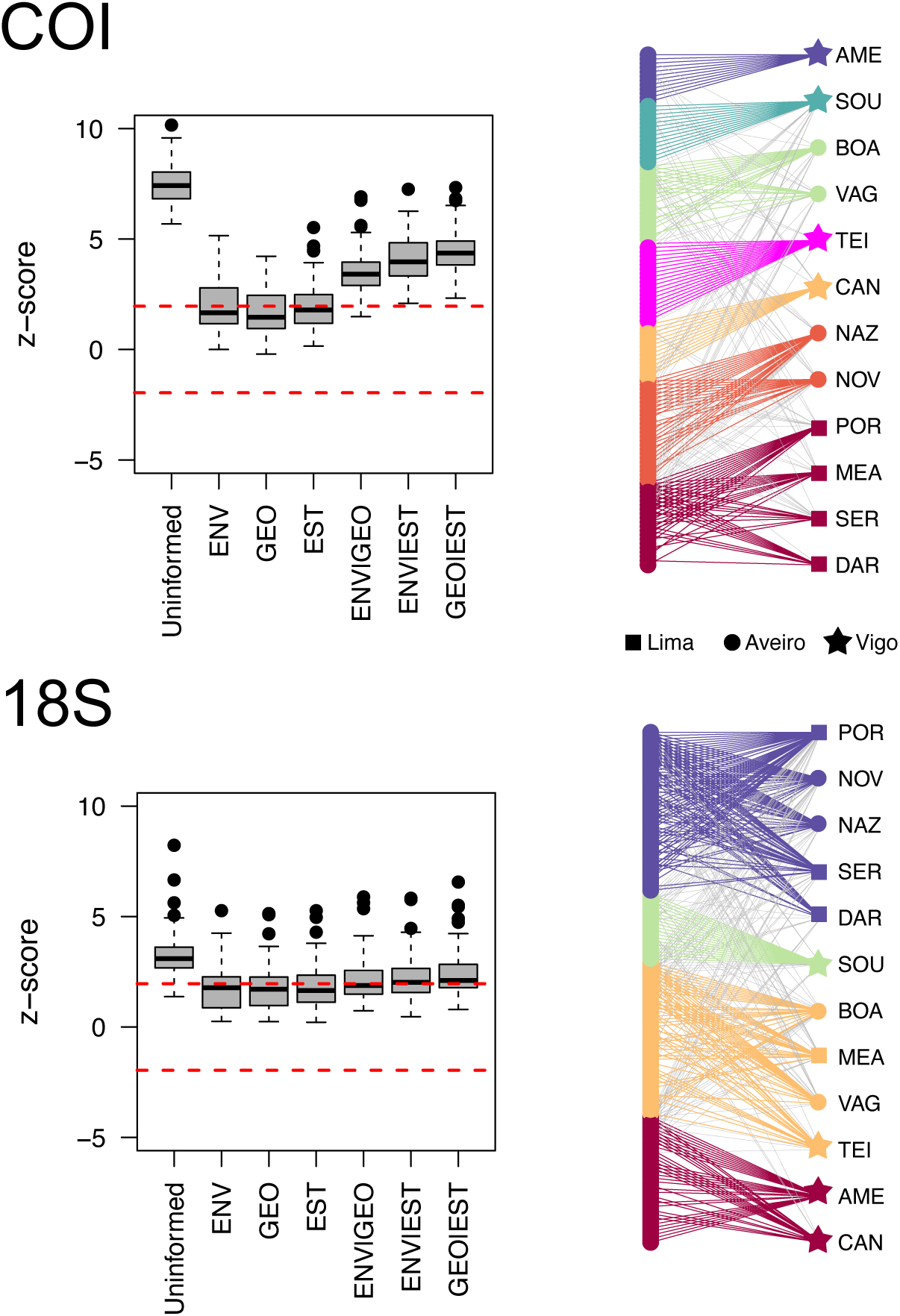
– Results from null model analyses and modular structures for the spatial networks surveyed by COI and 18S. Boxplots indicate different null model outcomes, with black circles indicating outliers and red dotted lines the thresholds for significance. Values of *z* ≤ -1.96 and *z* ≥ 1.96 indicate a significant under- and overrepresentation of the empirical modular structure, respectively. Acronyms in network graphs are for sampling stations (see Fig. 1 and Table 1), symbols for different estuaries and colours for identified modules. Coloured lines are for within module links and grey ones for between module links.

When considering the spatial network structure of meiofaunal OTUs sampled through 18S, we found a slightly lower modularity, *Q* = 0.373, with nodes distributed in four main modules. Differently from the modular partitioning observed for the COI network, we did not find a clear subdivision of sampling stations in well-defined modules (Fig. 3), with the only exception of three out of four located in Ría de Vigo, which belonged to different and mostly exclusive modules. None of informed null models were found to be particularly informative, meaning that neither the environmental differences nor the geographic positions between and within each estuary can be considered good predictors of the distributional pattern of meiofaunal communities sampled through the 18S.

### 3.4 Macroecological patterns of sampled OTUs

Factor Analysis of Mixed Data showed quite similar values (ca. 20%) of the total explained variance in both networks (Fig. 4). In the COI-derived networks, both the connectivity and participation coefficients were strongly correlated with the first axis (0.55 and 0.45, respectively) and only weakly correlated with the second axis (0.08 and 0.15, respectively). While the participation coefficient in the 18S-derived network showed a strong correlation with the first axis (0.56) and a weak correlation with the second axis (0.11), the connectivity coefficient showed quite similar correlations with both axes (0.30 and 0.33, respectively). The ordination of points in the multivariate space showed similar patterns for some phyla in the two spatial networks. This is the case for Amoebozoa, Annelida and Gastrotricha, which were characterized by relatively high values of both connectivity and participation coefficients, thereby having a widespread distribution encompassing different modules (Fig. 4). Conversely, Arthropoda Crustacea and Mollusca were characterized by lower values of connectivity and participation and, therefore, having a more restricted distributional range (Fig. 4). Other phyla, as Nematoda and Platyhelminthes showed opposite patterns of distribution, following positive values of connectivity and participation in the COI-derived network and negative in the 18S-derived networks (Fig. 4).

**Figure 4.**
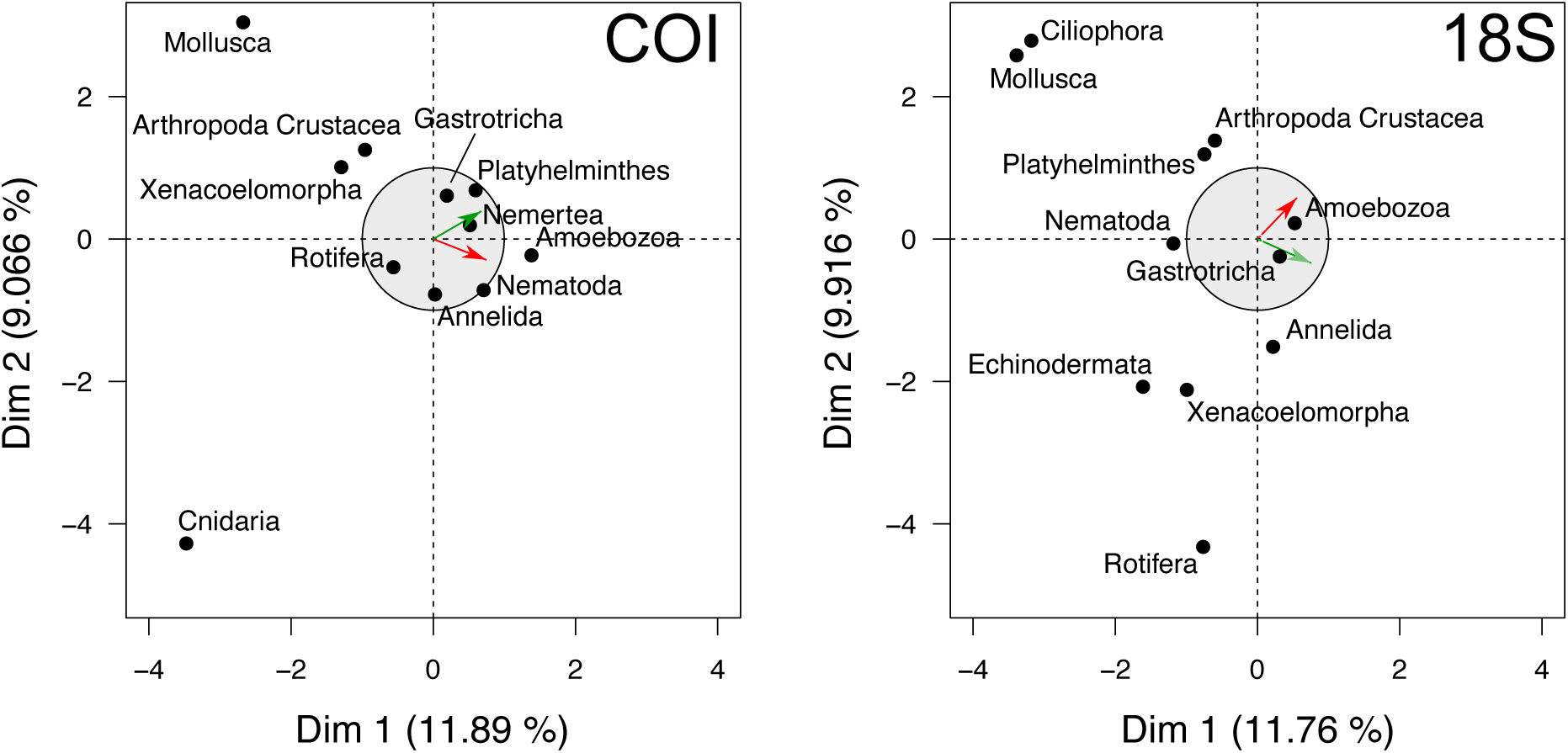
– Multivariate ordination pattern provided by the Factor Analysis of Mixed Data (FAMD) showing the distribution of phyla with respect to the connectivity (*z*, red arrow) and participation coefficients (*p*, green arrow). Light grey circles are the correlation circles (with radius of 1) depicting the proportion of variance captured by quantitative variables. The closer a variable is to the circle, the more important it is to interpreting the calculated principal dimensions. Variables on opposite sides of the origin are inversely correlated, whereas those on the same side are positively correlated.

## 4. Discussion

By integrating eDNA metabarcoding with network theory, we showed how the spatial arrangement of intertidal meiofaunal assemblages is largely modular, independently of the primers used. This suggests a non-random spatial aggregation of OTUs and sampling stations, meaning that both spatial networks show a significant subdivision in modules corresponding to geographical sub-regions representing the core area of current distribution of meiofaunal phyla within the system.

Modularity is a common property of many complex systems, arising as a natural emergent property in different types of networks. In spatial networks, modules identify critical meso- scales at which specific evolutionary and ecological processes act (Sadedin, 2005; Fortuna et al., 2008). For instance, previous studies have shown how movement and gene flow between populations cannot be explained by considering geographic distance alone, suggesting hidden spatial scales in metapopulation’s viability and the possibility that modules may constitute real evolutionary units (Fortuna et al., 2009). At the same time, modules may provide a fine assessment of the biogeographic boundaries in large-scale distributions of species, revealing the spatial limits at which dispersal and/or environmental filtering are involved in determining the pattern of species assemblage (Carstensen et al., 2012). Our findings are consistent with a modular subdivision which reflects fine scale differences between and within estuaries, with modules corresponding to local assemblages consisting of unique combinations of OTUs.

Mesoscale variability of subtidal meiobenthos has been previously reported (Alves et al., 2009), showing significant differences in the density and composition of meiofaunal assemblages at estuarine scale, but not over larger extent (i.e., between estuaries). Our findings not only confirm the high degree of patchiness in meiofaunal assemblages, but also show the ability of eDNA metabarcoding surveys to capture hierarchical spatial structures in the assemblage of intertidal meiofaunal assemblages. This pattern is particularly evident in the COI-derived spatial network, where OTUs and sampling stations aggregate in modules corresponding to estuaries or their subsets (Fig. 3). Indeed, all sampling stations in Lima belonged to the same module, while those of Ría de Aveiro were divided in two main modules corresponding to the most internal and external portions of the study area and those of Ría de Vigo to four distinct modules (Fig. 3). Such partition suggests heterogenous environmental conditions both within and between estuaries, probably related to the geomorphic characteristics of estuaries as, for instance, their position along the coast (e.g., parallel or perpendicular) or the degree of exposure with the sea, ultimately influencing the mixing of ocean and river waters and sedimentation processes (Fitzgerald et al., 2014).

By definition, estuaries are seaward portion of drowned valley systems formed during rising sea levels in the Pleistocene, being transition zones from freshwater entering through fluvial input and saline water from the open ocean (Pritchard, 1967; Fitzgerald et al., 2014). However, estuaries exist in a wide range of geological settings, including rias, glaciated valleys, structural basins and bar-built systems, each characterized by substantial differences in terms of stratification, hydrodynamics and sedimentation processes. The three estuary systems analysed in this study are all different according to Pritchard’s classification (1967). Ría de Vigo is classified as a fjord, thereby characterized by high length and depth, small width-to-depth ratios (ca. 1:10) and highly stratified waters (Fitzgerald et al., 2014); Lima is a drowned river valley characterized by mid-to-shallow water (ca. tens of metres), and a channel gradually widening and deepening towards the mouth (Fitzgerald et al., 2014); Ría de Aveiro is a bar-built estuary, characterized by shallow water (rarely exceeding 10 m in depth), large sediment carrying capacity and well-mixed waters (Fitzgerald et al., 2014).

Such a partition is less evident in the 18S-derived spatial network, where sampling stations do not follow a clear subdivision based on estuary affiliation (Fig. 3). Although the significant modular structure found in the network cannot be explained by local and regional differences in environmental features, this suggests complementary macroecological properties of meiofaunal communities identified by both primers, regardless of the type of monitored coastal ecosystem. These results are in line with previous studies in which different molecular markers were used for assessing microbial communities through a DNA metabarcoding/network analysis approach (Cobo-Diaz et al., 2019). Indeed, single-locus applications would lead not only to an inaccurate view of taxonomic and molecular diversity, but also to a lack of information on specific functional characteristics as, for instance, species response to local environmental conditions and differences in adaptations to interstitial life, unlikely to be detected without the integration of network analysis. Moreover, the combined use of DNA metabarcoding and network analysis would allow identifying the existence of recurrent and complementary macroecological characteristics of meiofaunal phyla with respect to the identified spatial structure and its determinants (Clare et al., 2019). Overall, our findings are consistent with a high degree of OTU turnover and ‘exclusivity’, also confirming the goodness of eDNA to detect macroecological patterns in meiofaunal communities.

Although our study focused on a limited part of community diversity, sample diversity within each station was successfully recovered, as confirmed by previous studies on meiofauna in Lima estuary (Fais et al., 2020a; 2020b). The present study not only re-captured the high turnover within and between the sampling stations in Lima estuary (Fais et al., 2020a; 2020b), but highlighted a similar situation in two other distinct estuarine systems in the region, spanning approx. 300 km maximum distance (Fig. 1). Detailed OTU sharing patterns comparison also suggests that exclusivity rather than pervasiveness appears to be the norm in meiofaunal organisms. In addition to the highly abundant nematodes, other taxonomic groups, such as platyhelminthes, gastrotriches, amoebozoans and cylophores, were also quite abundant among the intertidal meiofauna of different stations. Interestingly, the same phyla also show common macroecological features independently of the primers used, being characterized by high values of both connectivity and participation coefficients, thus having a widespread distribution encompassing different modules.

These results not only confirm the potential of metabarcoding in the analysis of meiofauna biodiversity at high taxonomic ranks (Faria et al., 2018; Leasi et al., 2018), but also the existence of a positive distribution-richness relationship in meiofaunal assemblages, hence identifying core (i.e., widespread and rich) and satellite (i.e., restricted and scarce) phyla (see Fig. 4 and Fig. S2 in Supplementary Materials). Overall, the results obtained by the integration of eDNA metabarcoding data and network analysis show that the combined use of COI and 18S is able to identify complementary taxonomic groups and macroecological features describing the pattern of diversity distribution in meiofaunal assemblages, ultimately emphasizing the importance of using a multilocus DNA metabarcoding approach (Clare et al., 2019).

## 5. Conclusions

Although the use of molecular data in network analysis is increasingly used with the aim to reconstruct species co-occurrence networks or even food webs, this study is the first, to our knowledge, to explicitly analyse meiofauna eDNA metabarcoding survey data within a spatial network framework. This allowed us to disentangle the role of environmental and geographic constraints on the distribution of meiofaunal assemblages in estuarine systems spanning different spatial scales, as well as the identification of common macroecological features of meiofaunal phyla.

Notwithstanding, actual limitations related to the correct identification of organisms at a specific level and/or the lack of reference libraries to improve the accuracy of taxonomic resolution in meiofauna might influence the outcomes of analyses. However, the use of a rigorous framework relating distributional data, environmental and geographic features, clearly demonstrates the feasibility of eDNA metabarcoding in recovering not only a fair amount of diversity signal at high taxonomic ranks (e.g., phyla), but also to identify recurrent spatial properties able to explain the pattern of diversity distribution. We argue that a similar approach, encompassing larger scales of investigation within and outside the Lusitanian province as well as the implementation of new bioinformatic procedures overcoming the limits of taxonomy-DNA based methodologies (e.g., supervised machine learning), might help to better understand the patterns and mechanisms of meiofauna distribution, shedding the light on the meiofauna paradox and helping to implement improved monitoring strategies.

## Supporting information

Supplementary results

## Acknowledgements

This work was supported by the Norte Portugal Regional Operational Programme (NORTE 2020) project ‘NextSea: Next generation monitoring of coastal ecosystems in a scenario of global change’ [NORTE-01-0145-FEDER-000032], under the PORTUGAL 2020 Partnership Agreement through the European Regional Development Fund (ERDF). MF and SD were also supported, respectively, by a PhD [SFRH/BD/113547/2015] and a postdoc [SFRH/BPD/109842/2015] fellowship provided by the Fundação para a Ciência e a Tecnologia (FCT). The authors would like to thank Prof. Jesús Troncoso (University of Vigo) and Prof. Ronaldo Sousa e Prof. Pedro Gomes (University of Minho) for their precious availability and collaboration over the years.

