## Supplementary results for "The spatial network structure of intertidal meiofaunal communities derived from environmental DNA metabarcoding surveys in Northwest Iberia"

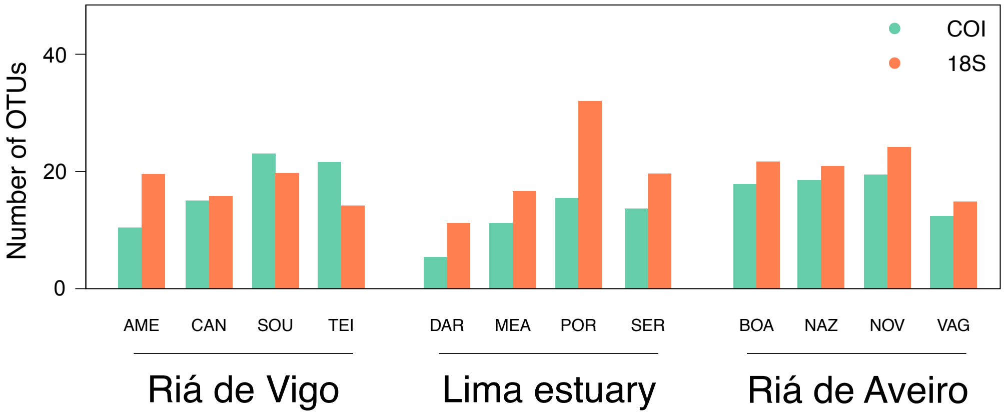


**Figure S1** – Barchart showing the number of OTUs surveyed by COI and 18S in different sampling stations and estuaries.

[1] Present address

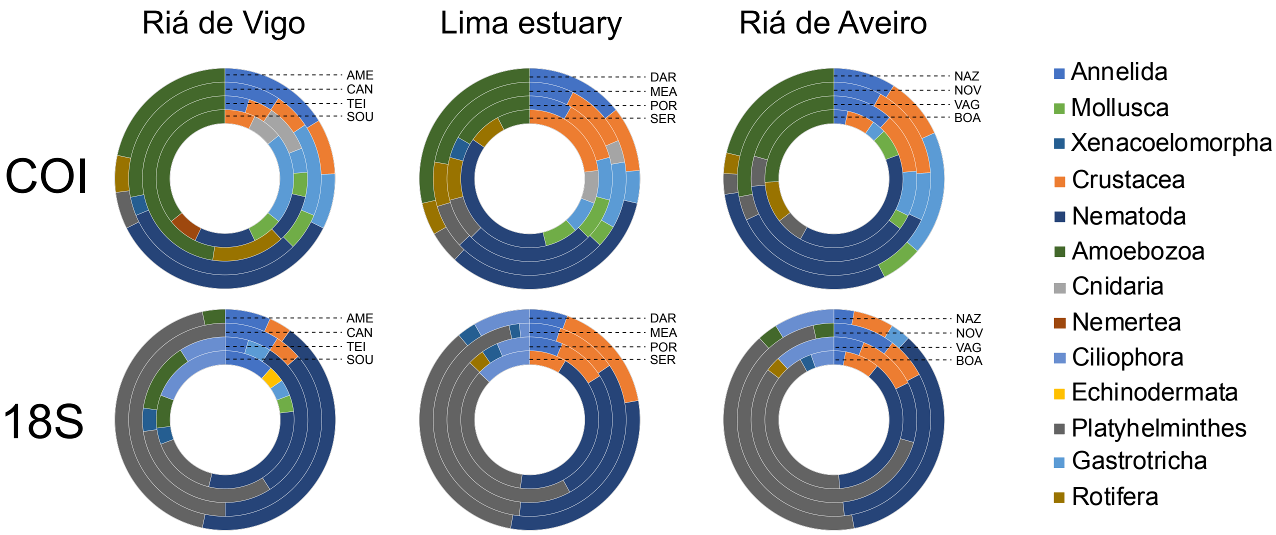


**Figure S2** – Doughnut charts showing the distribution of the main phyla in different sampling stations and estuaries.

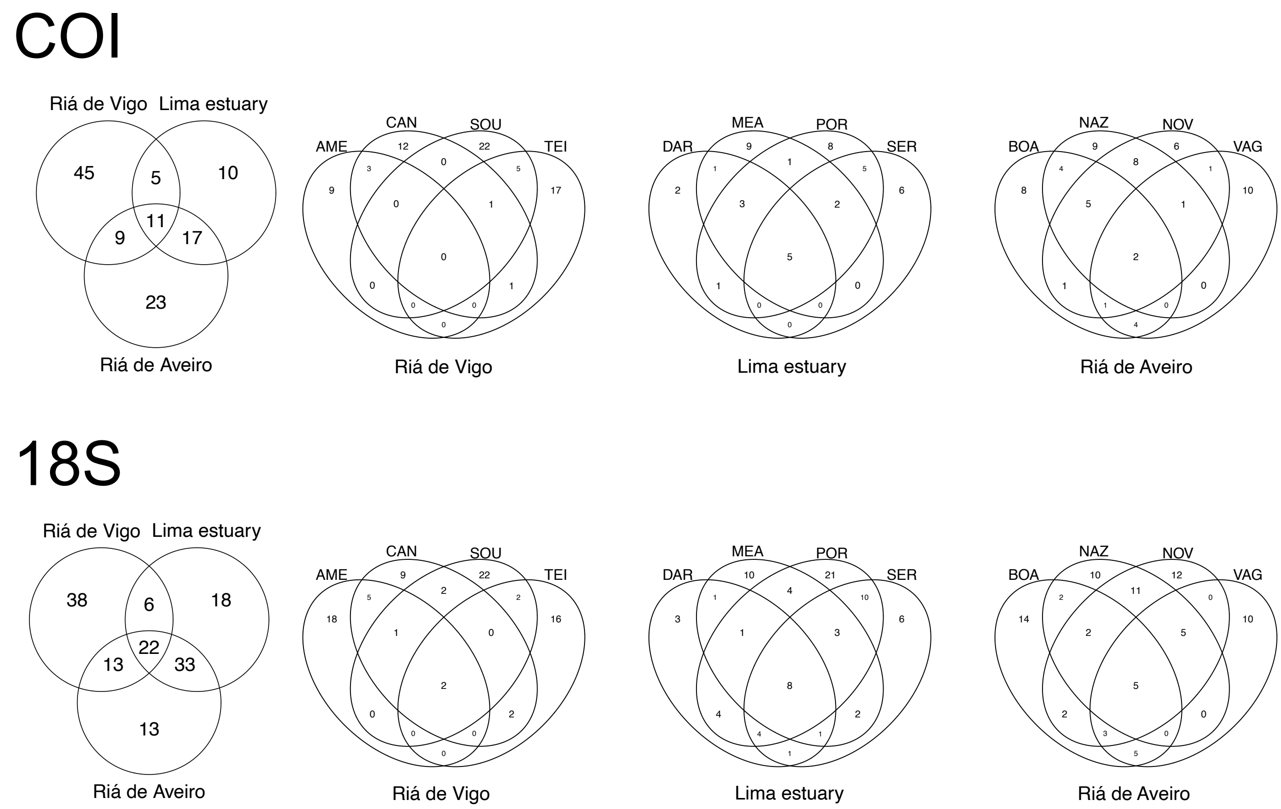


**Figure S3** – Venn diagram showing the shared pattern of OTUs between estuaries and between the sampling stations in each estuary.
